# Stalk Bending Strength is Strongly Associated with Maize Stalk Lodging Incidence Across Multiple Environments

**DOI:** 10.1101/675793

**Authors:** Rajandeep S. Sekhon, Chase N. Joyner, Arlyn J. Ackerman, Christopher S. McMahan, Douglas D. Cook, Daniel J. Robertson

## Abstract

Stalk lodging in maize results in substantial yield losses worldwide. These losses could be prevented through genetic improvement. However, breeding efforts and genetics studies are hindered by lack of a robust and economical phenotyping method for assessing stalk lodging resistance. A field-based phenotyping platform that induces failure patterns consistent with natural stalk lodging events and measures stalk bending strength in field-grown plants was recently developed. Here we examine the association between data gathered from this new phenotyping platform with counts of stalk lodging incidence on a select group of maize hybrids. For comparative purposes, we examine four additional predictive phenotypes commonly assumed to be related to stalk lodging resistance; namely, rind puncture resistance, cellulose, hemicellulose, and lignin. Historical counts of lodging incidence were gathered on 47 hybrids, grown in 98 distinct environments, spanning four years and 41 unique geographical locations in North America. Using Bayesian generalized linear mixed effects models, we show that stalk lodging incidence is associated with each of the five predictive phenotypes. Further, based on a joint analysis we demonstrate that, among the phenotypes considered, stalk bending strength measured by the new phenotyping platform is the most important predictive phenotype of naturally occurring stalk lodging incidence in maize, followed by rind puncture resistance and cellulose content. This study demonstrates that field-based measurements of stalk bending strength provide a reliable estimate of stalk lodging incidence. The stalk bending strength data acquired from the new phenotyping platform will be valuable for phenotypic selection in breeding programs and for generating mechanistic insights into the genetic regulation of stalk lodging resistance.

## 1. Introduction

Plant lodging, the permanent displacement of plants from their vertical stance (Carter and Hudelson, 1988; Rajkumara, 2008), causes major yield losses in several vital crop species including maize, wheat, and rice. Lodging is responsible for at least 5% annual yield loss in maize worldwide (Zuber and Kang, 1978; Flint-Garcia et al., 2003b). This estimate does not account for losses of maize stover that can be used in animal feed or biofuel feedstocks. Several factors are responsible for increased losses due to lodging in modern agriculture. In particular, cultivation of high yielding hybrids, application of high nitrogen, and increased planting density place increased mechanical demands on maize stalks (Duvick and Cassman, 1999). Moreover, improving the digestibility of lignocellulosic biomass to create better forage and biofuel feedstock is often associated with decreased stalk strength and hence increased incidence of stalk lodging (Pedersen et al., 2005; Feltus and Vandenbrink, 2012). Finally, an increase in number, frequency, and severity of extreme weather events resulting from climate change is expected to increase lodging incidence thus posing an ever-increasing production challenge in maize and other grasses.

### Box 1

**Description of key terms**

#### Lodging resistance

A conceptual, holistic assessment of the ability of a genotype to withstand external forces (wind and gravity) and the many biotic factors (insects, disease, etc.) that contribute lodging. This term refers to the overall behavior of a particular variety.

#### Predictive phenotypes

Phenotypic traits of a maize plant that are closely related to lodging resistance. The predictive phenotypes used in this study are defined below.

#### Bending strength

The measurement of the maximum amount of bending load that a stalk can tolerate. This is a per-plant measurement.

#### Lodging incidence

A count (or proportion) of plants lodged in a given area. This is a per-plot measurement.

#### Rind penetration resistance

The measurement of the maximum force required to press a thin rod through the rind of a stalk. This is per-plant measurement.

#### Lignin, Cellulose, and Hemicellulose Content

The proportion of dry weight of the stalk tissue that is composed of these structural macro-molecules. This may be a per-plant or per-plot measurement.

From an anatomical standpoint, lodging can either be caused by the inability of roots to keep the plant firmly implanted in soil, called root lodging, or by mechanical failure of the stalk (stem), termed stalk lodging. Root lodging in maize, characterized by leaning of the intact stalks at the crown root, is usually observed early during the season. Root lodging can cause a varying degree of yield losses (Bian et al., 2016). Stalk lodging, which involves breakage of a stalk below the ear, is usually observed as plants approach physiological maturity (Robertson et al., 2016) and often leads to total loss of the grain (Albrecht et al., 1986). Improving stalk lodging resistance (see Box 1 for description) offers an opportunity to substantially increase maize production without increasing agricultural inputs including irrigation, fertilizer, and crop management expenses.

Stalk lodging is a complex phenomenon determined by a complex interplay of a number of external and internal factors (Arnold and Josephson, 1975). Internal factors include metabolic and morphological properties of the plants that are expected to have genetic underpinnings. Metabolic properties refer to the amount and distribution of chemical compounds in the plant cell wall while morphological properties include the anatomical and geometrical features of the stalks. External factors include high-velocity winds, prolonged rain, and damage to maize stalks by certain insects (e.g., corn borers) and diseases (e.g., stem rot pathogens) (Smith and White, 1988; Flint-Garcia et al., 2003a; Quesada-Ocampo et al., 2016). In general, the relative contribution of each of these factors can vary both temporally (e.g., growing season) and spatially (e.g., planting zones or different anatomical regions within a single plant), thus challenging the development of a holistic understanding of the stalk lodging phenotype.

Deciphering the genetic architecture of stalk lodging resistance has been slow primarily due to lack of a reliable and reproducible phenotyping methodology to record this complex trait. Assessment of stalk lodging resistance has traditionally relied on late-season stalk lodging incidence counts obtained by recording the number of lodged plants in a defined area. Currently, lodging counts are the most widely used method for determining lodging resistance in breeding programs (Robertson et al., 2016). However, this method is confounded by numerous factors including varying levels of disease, pest pressure, soil fertility, wind velocity, and other weather and environmental conditions at the locations used for evaluation (Thompson, 1972; Hondroyianni et al., 2000; Flint-Garcia et al., 2003a; Hu et al., 2012; Robertson et al., 2016). Thus, it is necessary to acquire stalk lodging incidence data from multiple temporally and spatially distinct environments to get a reliable estimate of stalk lodging resistance of a genotype. These requirements impose a significant investment of time and resources for breeding programs. To alleviate this issue, several phenotypes that are believed to be related to stalk lodging, hereafter referred to as *<u>predictive phenotypes</u>*, have been considered as an alternative to counts of stalk lodging incidence.

Predictive phenotypes are generally related to either material or morphological properties of stalks that are believed to alter bending strength. Metabolic constituents of stalk cell walls including lignin, cellulose, and hemicellulose are considered to be important for imparting mechanical strength, but their association with stalk lodging is often debated (Davidson and Phillips, 1930; Appenzeller et al., 2004; Ching et al., 2006). Furthermore, collecting data on these phenotypes is laborious, time consuming, and expensive. Rind puncture resistance recorded by an electronic rind penetrometer is another commonly used predictive phenotype (Sibale et al., 1992). Rind puncture resistance shows significant negative correlations with natural occurrence of stalk lodging in the field and provides higher sample throughput than metabolic analyses (Hu et al., 2012). However, rind puncture resistance is a one-dimensional variable that lacks inclusiveness of many biomechanical, metabolic, and morphological indicators of stalk lodging incidence. Various measurements of stalk strength have also been shown to be related to stalk lodging incidence in maize (Zuber and Grogan, 1961; Singh, 1970; Hu et al., 2013; Ma et al., 2014; Robertson et al., 2014; Robertson et al., 2016). However, the available strength measurement techniques require transporting stalks to a laboratory and the use of expensive material testing equipment. Most breeding programs have, therefore, found measurements of stalk strength to be unsuitable for breeding purposes due to a high investment of time, labor, and cost.

To address issues that circumvent adoption of stalk strength as a predictive phenotype for stalk lodging incidence, a novel field-based platform for assessing stalk bending strength of maize stalks *in vivo* was developed. The novel device, known as DARLING (Device for Assessing Resistance to Lodging IN Grains), measures stalk bending strength in field-grown plants by simulating the loading conditions and failure patterns of naturally lodged plants (Cook et al., 2019). Stalk bending strength data acquired from the device are strongly correlated with lab-based measurements of stalk bending strength and stalk flexural stiffness (Cook et al., 2019). Stalk bending strength is determined by the cumulative effect of metabolic and morphological properties of stalks. However, the extent to which the stalk bending strength measurements produced by DARLING are related to stalk lodging incidence in field environments has not been investigated.

The main objectives of this study are 1) to evaluate the association between field-based stalk bending strength data obtained by DARLING and counts of stalk lodging incidence, and 2) to compare the DARLING method with more traditional predictive phenotyping methods for stalk lodging incidence. Findings from this study will help in establishing a robust phenotyping strategy, boosting efforts for genetic dissection and breeding of stalk lodging resistance, and enhancing maize production.

## 2. Materials and methods

### 2.1. Acquisition of stalk lodging incidence data

In the current study, forty-seven maize hybrids were evaluated for stalk lodging incidence, and for five predictive phenotypes including stalk bending strength, rind puncture resistance, cellulose, hemicellulose, and lignin (Table 1). Comprehensive data on naturally occurring stalk lodging incidence in a select group of maize hybrids was obtained from the Genome to Fields (G2F) initiative (www.genomes2fields.org). The G2F initiative is a multi-institutional collaboration with a goal of developing and improving tools to predict phenotypic performance of diverse maize genotypes across multiple growing conditions (AlKhalifah et al., 2018). In the G2F initiative, data collected by each of the G2F collaborators are submitted to a shared data repository and curated by a core team led by G2F project leaders.

**Table 1:**
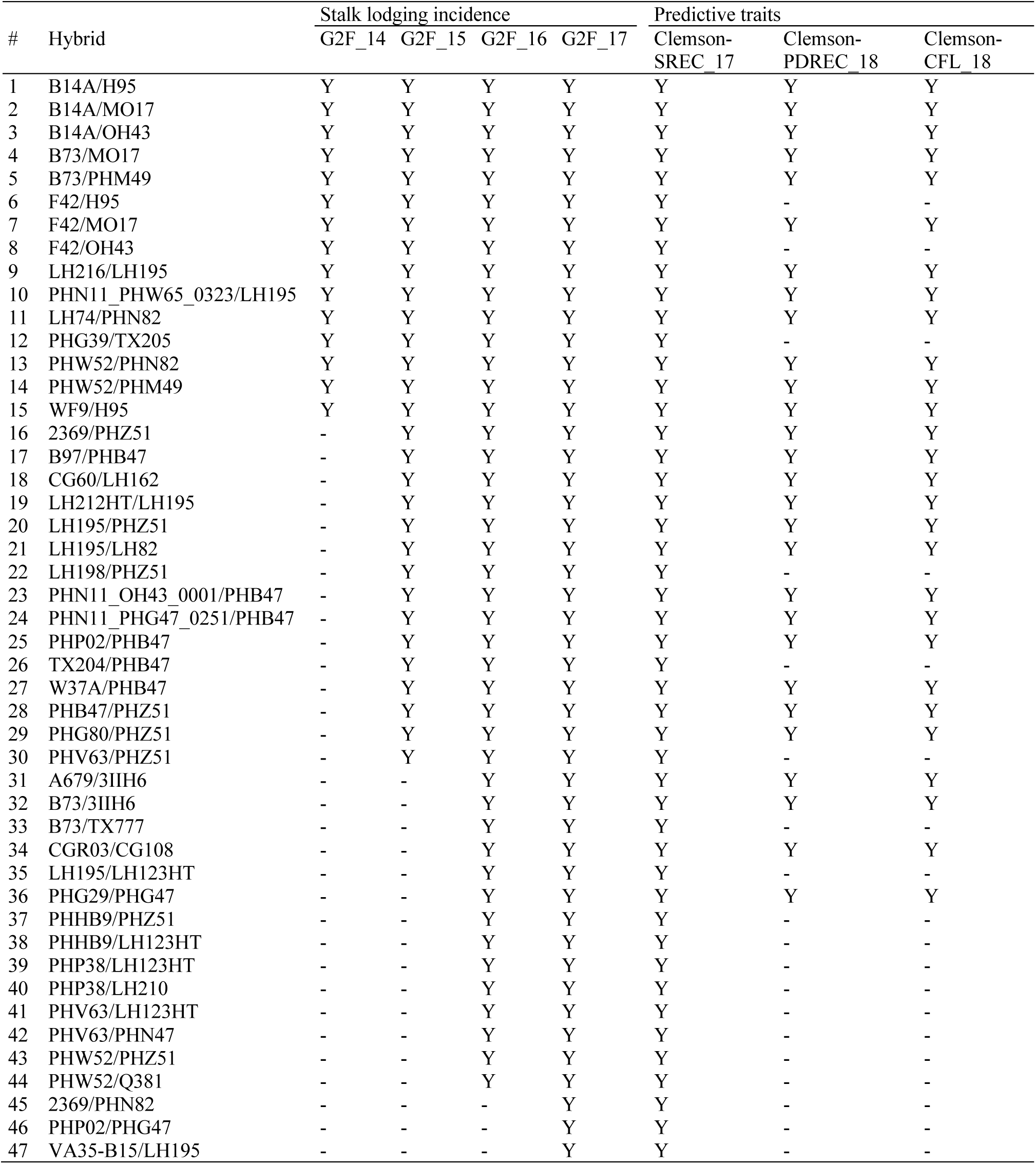
Details of hybrids and environments used in the study

| # | Hybrid | Stalk lodging incidence |  |  |  | Predictive traits |  |  |
| --- | --- | --- | --- | --- | --- | --- | --- | --- |
|  |  | G2F_14 | G2F_15 | G2F_16 | G2F_17 | Clemson-SREC 17 | Clemson-PDREC 18 | Clemson-CFL 18 |
| 1 | B14A/H95 | Y | Y | Y | Y | Y | Y | Y |
| 2 | B14A/MO17 | Y | Y | Y | Y | Y | Y | Y |
| 3 | B14A/OH43 | Y | Y | Y | Y | Y | Y | Y |
| 4 | B73/MO17 | Y | Y | Y | Y | Y | Y | Y |
| 5 | B73/PHM49 | Y | Y | Y | Y | Y | Y | Y |
| 6 | F42/H95 | Y | Y | Y | Y | Y | - | - |
| 7 | F42/MO17 | Y | Y | Y | Y | Y | Y | Y |
| 8 | F42/OH43 | Y | Y | Y | Y | Y | - | - |
| 9 | LH216/LH195 | Y | Y | Y | Y | Y | Y | Y |
| 10 | PHN11_PHW65_0323/LH195 | Y | Y | Y | Y | Y | Y | Y |
| 11 | LH74/PHN82 | Y | Y | Y | Y | Y | Y | Y |
| 12 | PHG39/TX205 | Y | Y | Y | Y | Y | - | - |
| 13 | PHW52/PHN82 | Y | Y | Y | Y | Y | Y | Y |
| 14 | PHW52/PHM49 | Y | Y | Y | Y | Y | Y | Y |
| 15 | WF9/H95 | Y | Y | Y | Y | Y | Y | Y |
| 16 | 2369/PHZ51 | - | Y | Y | Y | Y | Y | Y |
| 17 | B97/PHB47 | - | Y | Y | Y | Y | Y | Y |
| 18 | CG60/LH162 | - | Y | Y | Y | Y | Y | Y |
| 19 | LH212HT/LH195 | - | Y | Y | Y | Y | Y | Y |
| 20 | LH195/PHZ51 | - | Y | Y | Y | Y | Y | Y |
| 21 | LH195/LH82 | - | Y | Y | Y | Y | Y | Y |
| 22 | LH198/PHZ51 | - | Y | Y | Y | Y | - | - |
| 23 | PHN11_OH43_0001/PHB47 | - | Y | Y | Y | Y | Y | Y |
| 24 | PHN11_PHG47_0251/PHB47 | - | Y | Y | Y | Y | Y | Y |
| 25 | PHP02/PHB47 | - | Y | Y | Y | Y | Y | Y |
| 26 | TX204/PHB47 | - | Y | Y | Y | Y | - | - |
| 27 | W37A/PHB47 | - | Y | Y | Y | Y | Y | Y |
| 28 | PHB47/PHZ51 | - | Y | Y | Y | Y | Y | Y |
| 29 | PHG80/PHZ51 | - | Y | Y | Y | Y | Y | Y |
| 30 | PHV63/PHZ51 | - | Y | Y | Y | Y | - | - |
| 31 | A679/3IIH6 | - | - | Y | Y | Y | Y | Y |
| 32 | B73/3IIH6 | - | - | Y | Y | Y | Y | Y |
| 33 | B73/TX777 | - | - | Y | Y | Y | - | - |
| 34 | CGR03/CG108 | - | - | Y | Y | Y | Y | Y |
| 35 | LH195/LH123HT | - | - | Y | Y | Y | - | - |
| 36 | PHG29/PHG47 | - | - | Y | Y | Y | Y | Y |
| 37 | PHHB9/PHZ51 | - | - | Y | Y | Y | - | - |
| 38 | PHHB9/LH123HT | - | - | Y | Y | Y | - | - |
| 39 | PHP38/LH123HT | - | - | Y | Y | Y | - | - |
| 40 | PHP38/LH210 | - | - | Y | Y | Y | - | - |
| 41 | PHV63/LH123HT | - | - | Y | Y | Y | - | - |
| 42 | PHV63/PHN47 | - | - | Y | Y | Y | - | - |
| 43 | PHW52/PHZ51 | - | - | Y | Y | Y | - | - |
| 44 | PHW52/Q381 | - | - | Y | Y | Y | - | - |
| 45 | 2369/PHN82 | - | - | - | Y | Y | - | - |
| 46 | PHP02/PHG47 | - | - | - | Y | Y | - | - |
| 47 | VA35-B15/LH195 | - | - | - | Y | Y | - | - |

The lodging incidence data was based on visual observation made at physiological maturity. Counts of stalk lodging incidence were obtained from 98 distinct G2F environments spanning four years (2014, 2015, 2016, and 2017), 41 unique geographical locations across 19 states in the US, and one province in Canada, (Table B.1.). Geographically, these environments ranged from latitudes between 30.55° and 65.30° and longitudes between -105.00° and -75.20°. Details of specific growing conditions and agronomic practices for these environments are available in supplementary Table B.1. While this dataset included stalk lodging incidence for 47 hybrids, not every hybrid was tested in all environments. The number of test environments for a hybrid ranged from 17 to 93 with an average of 55 environments.

### 2.2. Experimental design for assessment of predictive phenotypes

Predictive phenotypes including bending strength, rind puncture resistance, and metabolic constituents (cellulose, hemicellulose, and lignin) were evaluated in three environments spanning two years (2017 and 2018) and three unique geographical locations (Table 1). In 2017, the hybrids of interest were planted at Clemson University (CU) Simpson Research and Education Center (Clemson-SREC), Pendleton, SC. The soil in Clemson-SREC is 100% well drained Cecil sandy loam, consisting of sandy loam in the top 0-6 inches and mostly clay beneath this top layer. In 2018, the same hybrids were planted at CU Calhoun Field Laboratory (Clemson-CFL) and at CU Pee Dee Research and Education Center (Clemson-PDREC), Florence, SC. The soil type in Clemson-CFL is Toccoa soil sandy loam to fine sandy loam. The area lays in a floodplain of Lake Hartwell and, therefore, moisture levels were consistent at field capacity for the entirety of the growing season, reaching saturation levels on occasional heavy rains. The soil at Clemson-PDREC is Norfolk loamy sand. At all three environments, the hybrids were grown in a Random Complete Block Design with two replications. In each replication, each hybrid was planted in two-row plots (4.57 m length and 0.76 m apart) at a planting density of 70,000 plant ha^-1^. For each experiment, 56.7 kg ha^-1^ nitrogen was added at soil preparation and additional 85 kg ha^-1^ nitrogen applied 30 days after emergence. Standard agronomic practices were followed for crop management. In 2017, data were collected on 47 hybrids from Clemson-SREC environment and, in 2018, data were collected on 28 hybrids from Clemson-CFL and Clemson-PDREC environments (Table 1).

### 2.3. Sampling strategy for predictive phenotypes

Bending strength data, rind puncture data, and metabolic data were collected at physiological maturity when all the hybrids were either at or past 40 days after anthesis. Data were collected on 10 randomly chosen competitive plants in each plot. Nominally, 60 total measurements were acquired for each hybrid (10 plants per plot × 2 replications × 3 locations). However, some plots lacked 10 competitive plants and, therefore, the total number of plants phenotyped for each hybrid varied slightly. Rind puncture tests were performed on the same set of plants that were tested in bending. Metabolic data were collected on all 47 hybrids grown in Clemson-SREC environment in 2017. Samples for metabolic data were acquired from the same plants that were subjected to bending tests and rind puncture tests.

### 2.4. Stalk bending strength methodology

Stalk bending strength was measured using the newly developed Device for Assessing Resistance to Lodging IN Grains (DARLING). A detailed description and proper usage of DARLING and the type of data collected has been published recently (Cook et al., 2019). Briefly, the DARLING apparatus consists of a vertical arm and a footplate. A force sensor mounted on the vertical arm is aligned with the designated stalk by placing the footplate flush with the base of the stalk. The force sensor is placed at the center of the internode beneath the ear and, by pushing it forward, the device rotates on a fulcrum at the intersection of the vertical arm and footplate to push the stalk over (Fig. 1). The force applied to the stalk induces a bending moment in the stalk similar to the bending moment created by wind loading and creates a failure pattern that is consistent with natural lodging (Robertson et al., 2015; Cook et al., 2019).

**Fig. 1.**
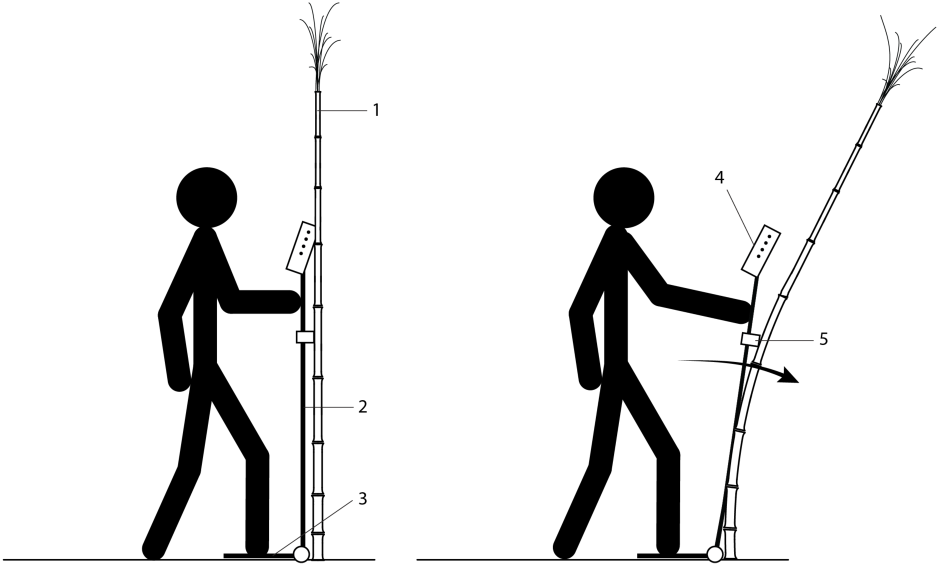
Illustration of the manner in which DARLING is used to test maize stalks.

While being pushed forward by the user, the device continuously measures the deflected angle using a potentiometer. The force sensor likewise continuously collects data from the resistive force exerted by the stalk. The resulting force/rotation chart is shown to the user using an onboard screen. A smooth push of a stalk with correct placement of the load cell results in data points arcing upward until stalk failure occurs. At the point of stalk failure, a distinct drop in force is observed with continued deflection. At this point, the DARLING apparatus records the maximum bending moment exerted on the stalk prior to failure and the load cell is reset for the next measurement. Since failure can be due to stalk breakage, root lodging, or a combination of the two, the user enters a note after each measurement specifying the nature of the failure.

### 2.5. Rind puncture resistance methodology

Rind puncture resistance was measured with an Imada® Digitial Force Gauge (ZTA-DPU) outfitted with a flat tipped probe of 2 mm diameter (item #49FL81, Grainger Inc., Lake Forest, IL). The digital force gauge was modified with a steel plate covering the base of the probe to prevent secondary contact post-puncture. The internode below the primary ear-bearing node was used for rind puncture testing after careful removal of the leaf sheath. The location of the rind puncture was in the center of the internode, perpendicular with the minor axis, and in the direction of the major axis. To eliminate inter-user variability, the same single user conducted puncture testing for all samples included in the study.

### 2.6. Metabolic analysis methodology

Three internodes below the primary ear-bearing node were harvested along with the intervening nodes for metabolic analysis. The internodes were dried down in a forced air drier at 65°C. Dried internodes from a plot were bulked and ground in SM 300 cutting mill (Retsch GmbH, Germany). For each plot, 25 grams of ground tissue was submitted to Dairyland Laboratories, Inc. (Arcadia, WI) for wet chemistry analysis. Neutral detergent fiber (NDF), Acid detergent fibers (ADF), and lignin values were measured using the standardized AOAC official methods. Cellulose values were obtained by calculating the difference between ADF and lignin content of samples. Hemicellulose values were obtained by calculating the difference between NDF and ADF of samples.

### 2.7. Quality control and preparation of data for analysis

The focus of this study was to examine the significance and determine variable importance of several predictive phenotypes with regard to their association with stalk lodging incidence. As such, any plants that did not experience stalk breakage during testing with DARLING were excluded from the data set. In other words, the data from plants that root lodged when tested by the DARLING apparatus were discarded. However, since metabolic data were generated on a pooled sample from 10 plants in a plot, these data are from plants that showed all types of failure patterns during testing with DARLING. Lastly, any extreme outlying measurements (i.e., bending strength measurements < 0.1 Nm and rind puncture resistance measurements greater than 40 lbf) were removed prior to analysis, as these data approach or exceed the limits of the phenotyping devices employed in the study.

### 2.8 Statistics

We relate counts of stalk lodging incidence to predictive phenotypes (stalk bending strength, rind puncture resistance, cellulose, hemicellulose, and lignin) through two Bayesian generalized linear mixed effects models. Mixed-effects models are common in crop ecology where the data often contains clusters (i.e., fields) of observational units (Zeger and Karim, 1991; Bolker et al., 2009; Nakagawa and Schielzeth, 2013). Let *s*_*ij*_, *r*_*ij*_, *c*_*ij*_, *h*_*ij*_, and *l*_*ij*_ denote stalk bending strength, rind puncture resistance, cellulose, hemicellulose, and lignin, respectively, for the *i*th plant at the *j*th environment. For comparative purposes, all predictive phenotypes were standardized; i.e., proceeding in this fashion allows one to assess variable importance by simply examining the estimated regression coefficients, with larger (in magnitude) estimates relating to higher degrees of importance. To account for between-field variation due to unknown environmental factors, a random intercept *γ*_*j*_ ∼ *N*(0, *τ*^2^) is shared for all plants at the *j*th environment. For completeness, we consider both a marginal and a joint model that are given by

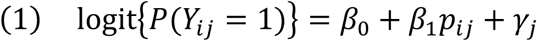

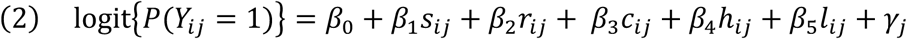

respectively, where *Y*_*ij*_ = 1 denotes the event that the *i*th plant in the *j*th field lodged, and *Y*_*ij*_ = 0 otherwise, for *i* = 1, …, *n*_*j*_ and *j* = 1, …, 98. In (1), we use *p*_*ij*_ to arbitrarily denote the five predictive phenotypes; i.e., *s*_*ij*_, *r*_*ij*_, *c*_*ij*_, *h*_*ij*_, and *l*_*ij*_. A few comments are warranted. First, the usual conditioning arguments in the probability statements have been dropped in models (1) and (2) for notional brevity. Second, model (1) is a marginal model with a random intercept which we fit using each of the predictive phenotypes one at a time while model (2) is a joint model which considers all of the predictive phenotypes simultaneously. It is important to note that stalk lodging incidence and the predictive phenotypes were collected in separate experiments. In particular, lodging incidence was gathered from 98 environments spanning four years (2014, 2015, 2016, and 2017) and 41 unique geographical locations, whereas predictive phenotypes were acquired at three environments across two years (2017 and 2018). Thus, to harmonize the data, we used regression imputation to impute the predictive phenotypes for the stalk lodging incidence data (Lokupitiya et al., 2006). In particular, we train a predictive model using the data captured by the techniques outlined above with genotype being the only explanatory factor. Proceeding in this fashion, we impute predictive phenotypes by genotype for the G2F data as the average of the observed. For example, if 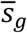 denotes the average of all available surrogate strength measurements for the *gth* maize variety, then set 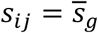 if the *i*th plant in the *j*th field is of the *g*th maize variety.

Model fitting for (1) and (2) was completed via Markov chain Monte Carlo (Hoff, 2009; Brooks et al., 2011). Convergence diagnostics, parameter estimation, and inference proceeded in the usual manner (Hoff, 2009; Brooks et al., 2011). Tables 2 and 3 provide a summary of the parameter estimates under models (1) and (2), respectively. These results include point estimates of regression parameters (i.e., estimates of the posterior mean) and estimates of 95% credible intervals, the Bayesian counterpart of the usual confidence intervals (Hoff, 2009). All estimates are based on 2,000 Markov chain Monte Carlo posterior draws that were retained after a sufficient burn-in period.

**Table 2:** Summary of the parameter estimates obtained from the marginal model. Included are the point estimates (Estimate) of the parameters (i.e., posterior mean) and associated measures of uncertainty (i.e., estimated standard deviation of the posterior distribution, SD). Also included are 95% Credible Intervals (CI). Note, significance can be ascertained by examining whether the CI’s contain zero.

| Trait | Parameter | Estimate | SD | 95% CI |
| --- | --- | --- | --- | --- |
| Stalk bending strength | Intercept | -3.91 | 0.09 | -4.07, -3.75 |
|  | Effect | -0.38 | 0.01 | -0.39, -0.36 |
| Rind puncture resistance | Intercept | -4.07 | 0.09 | -4.22, -3.90 |
|  | Effect | -0.30 | 0.01 | -0.32, -0.29 |
| Cellulose | Intercept | -4.23 | 0.13 | -4.48, -4.04 |
|  | Effect | 0.17 | 0.01 | 0.15, 0.19 |
| Hemicellulose | Intercept | -3.92 | 0.06 | -4.04, -3.81 |
|  | Effect | -0.13 | 0.01 | -0.14, -0.11 |
| Lignin | Intercept | -3.78 | 0.07 | -3.90, -3.67 |
|  | Effect | 0.24 | 0.01 | 0.22, 0.25 |

**Table 3:** Summary of the parameter estimates obtained from the joint model. Included are the point estimates (Estimate) of the parameters (i.e., posterior mean) and associated measures of uncertainty (i.e., estimated standard deviation of the posterior distribution, SD). Also included are 95% Credible Intervals (CI). Note, significance can be ascertained by examining whether the CI’s contain zero.

| Parameter/Trait | Estimate | SD | 95% CI |
| --- | --- | --- | --- |
| Intercept | -3.77 | 0.11 | -3.94, -3.58 |
| Stalk bending strength | -0.38 | 0.01 | -0.40, -0.36 |
| Rind puncture resistance | -0.13 | 0.01 | -0.15, -0.11 |
| Cellulose | -0.29 | 0.01 | -0.32, -0.26 |
| Hemicellulose | 0.06 | 0.01 | 0.05, 0.08 |
| Lignin | 0.36 | 0.01 | 0.34, 0.39 |

## 3. Results

### 3.1 Variation for stalk lodging incidence

Stalk lodging incidence data showed substantial variation across 98 environments spanning 19 US states and a Canadian province of Ontario over a period of four years (Fig. 2). The number of plants per plot ranged from 5 to 129 with an average of 61 and a median of 63 plants (Fig. 2A). The number of lodged plants in a plot ranged from 0 to 87 with an average of 3.7 and a median of 0 (Fig. 2B). Over the four years, overall stalk lodging incidence was the lowest in 2015 (4.3%) and the highest in 2016 (8%) (Fig. 2C). Overall stalk lodging incidence in each environment ranged between 0% and 46%. The states with the lowest and highest degree of stalk lodging incidence were Arkansas (0.5%) and Colorado (19.7%), respectively. Stratifying by time, Fig. 2 depicts stalk lodging incidence for each state by year. Further parsing the data, Fig. 3 provides the stalk lodging incidence after stratifying by hybrid type and environment; presented are box plots of the different incidences observed for each hybrid across the different environments. These results indicate a sizeable variation in stalk lodging incidence that is attributable to both the hybrid type and the environment. In fact, the plot level stalk lodging incidence of individual hybrids across all environments ranged between 0 and 100% with an average of 6% and a median of 0%.

**Fig. 2.**
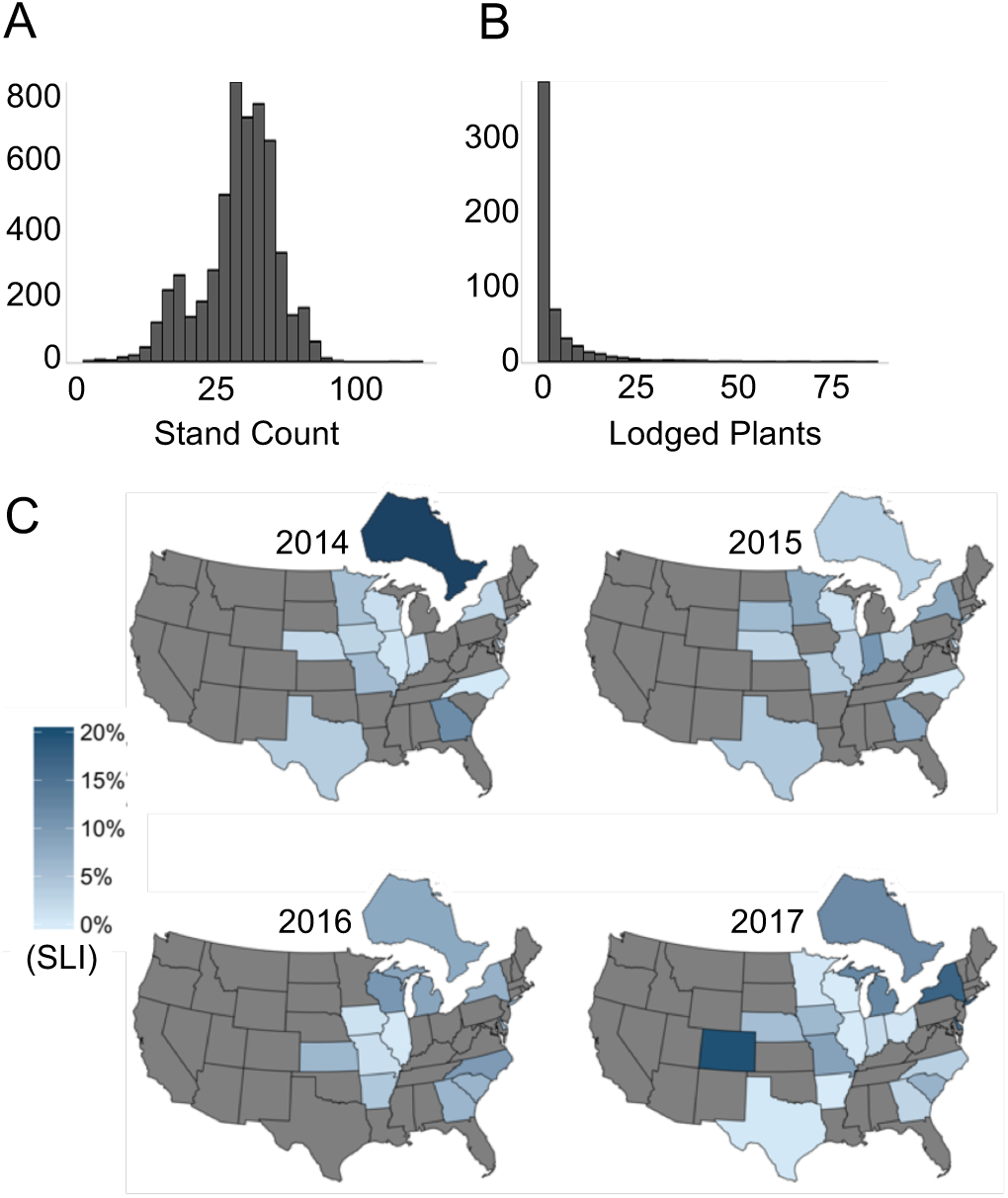
Spatial and temporal variation for stalk lodging incidence in the Genomes to Fields (G2F) initiative. (A) Frequency distribution of the total number of plants in a plot (stand count) across all locations and years, (B) The number of lodged plants in a plot across all locations and years, and (C) The extent of lodging in 19 US states and one Canadian province. For each year, shading of a state/province is based on the overall stalk lodging incidence across all considered hybrids at G2F locations in the state/province. States not reporting data are shaded gray.

**Fig. 3.**
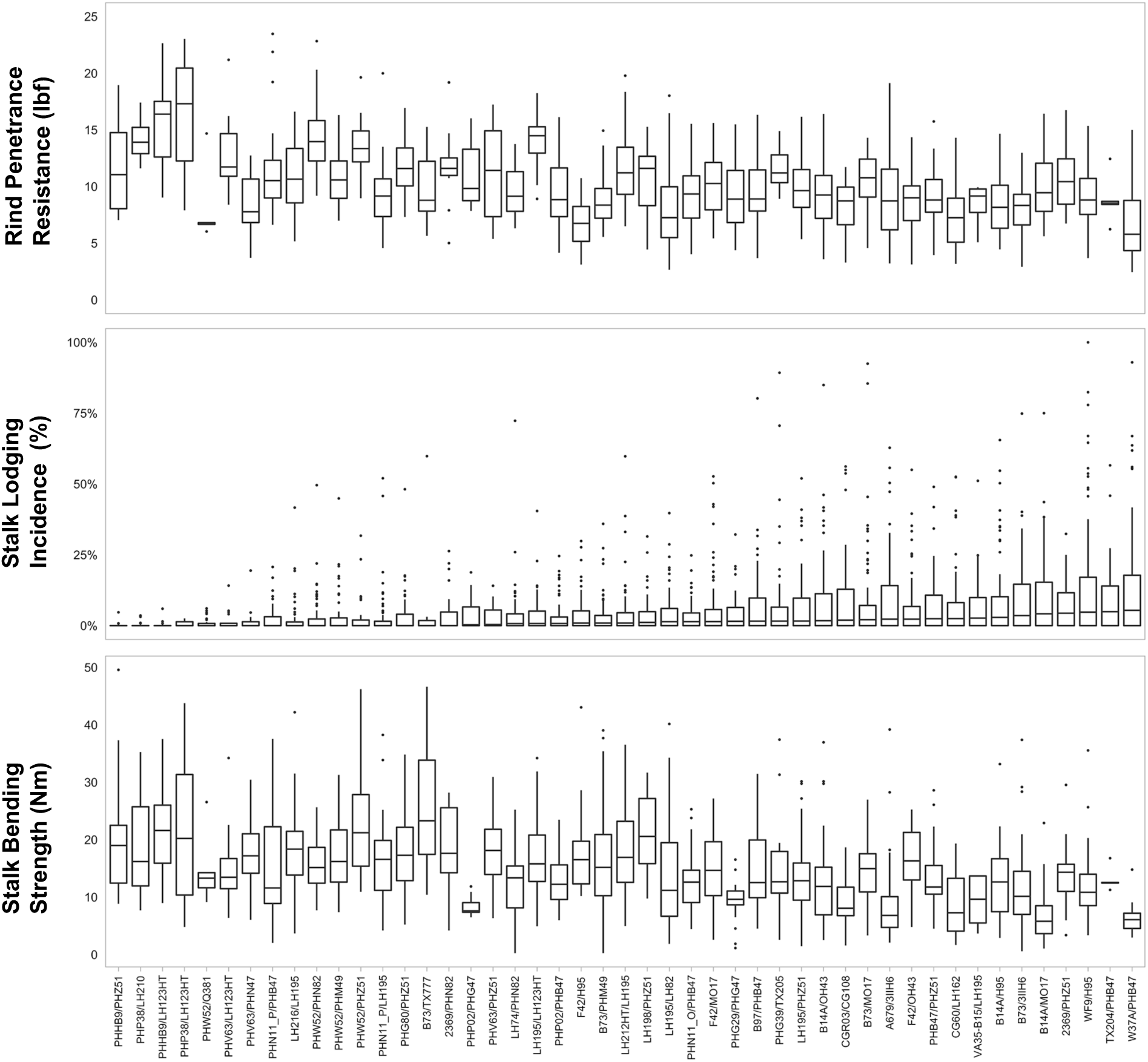
Variability in stalk lodging incidence and related traits captured in the hybrids included in the study. The hybrids are arranged based on increasing median stalk lodging incidence (middle panel). Box plots show the empirical distribution of three phenotypes; RPR (top panel), stalk lodging incidence (middle panel), and stalk bending strength (bottom panel). In the box plots the lower and upper end of the boxes represent 25^th^ and 75^th^ percentiles of the observed data, respectively; the tips of the vertical lines (whiskers) at the lower and upper end of the boxes represent the 10^th^ and the 90^th^ percentiles of the observed data, respectively; horizontal line within each box denotes the median, and the dots represent outliers. The following hybrid names were shortened: PHN11_PHG47_0251/PHB47 to PHN11_P/PHB47, PHN11_PHW65_0323/LH195 to PHN11_P/LH195, and PHN11_OH43_0001/PHB47 to PHN11_O/PHB47.

### 3.2 Variation in predictive phenotypes

To compare the relative importance of the predictive phenotypes with regard to their association with stalk lodging incidence, the predictive phenotypes were recorded for the same 47 hybrids mentioned above in three distinct environments, spanning two years. Measurements of stalk bending strength with DARLING revealed substantial differences among hybrids in all the environments (Fig. 3, supplementary Fig. A.1.) and the stalk bending strength of individual hybrids ranged from 0.25 to 49.56 Nm with an average of 14.22 Nm and a median of 12.99 Nm. Likewise, considerable variation was observed for rind puncture resistance in these hybrids ranging from 2.45 to 23.47 lbf with a mean of 9.96 lbf and a median of 9.56 lbf (Fig. 3, supplementary Fig. A.1.).

Since material properties of the stalks are also considered to be important to strength, we measured cellulose, hemicellulose, and lignin as percent of the dry weight of stalks. Cellulose content ranged from 33.81 to 48.58% with an average of 41.86% and a median of 42.65% (supplementary Fig. A.2.). Low variation was observed for hemicellulose content which ranged from 25.2 to 33.4% with an average of 31.04% and a median of 31.29%. Finally, lignin content of the hybrids ranged from 6.75 to 9.56% with an average of 8.02% and a median of 8%.

### 3.3 Association between stalk lodging incidence and the predictive phenotypes

We examined the relationship between each predictive phenotype and stalk lodging incidence using both of the aforementioned Bayesian generalized linear mixed effects models. We first tested the association between each of the predictive phenotypes and stalk lodging incidence individually. This analysis revealed that the linear term (effect) associated with each of the predictive phenotypes was significant at the 0.05 significance level (Table 2, Fig. 4). The stalk bending strength measurements produced by DARLING had the largest (in magnitude) effect estimate among all phenotypes and was negative. This indicates that an increase in stalk bending strength is associated with a decrease in stalk lodging incidence. Rind puncture resistance and hemicellulose also showed a negative association with stalk lodging incidence. Lignin had the largest positive effect estimate followed by cellulose, indicating a positive association between these metabolites and stalk lodging incidence, which was not expected.

**Fig. 4.**
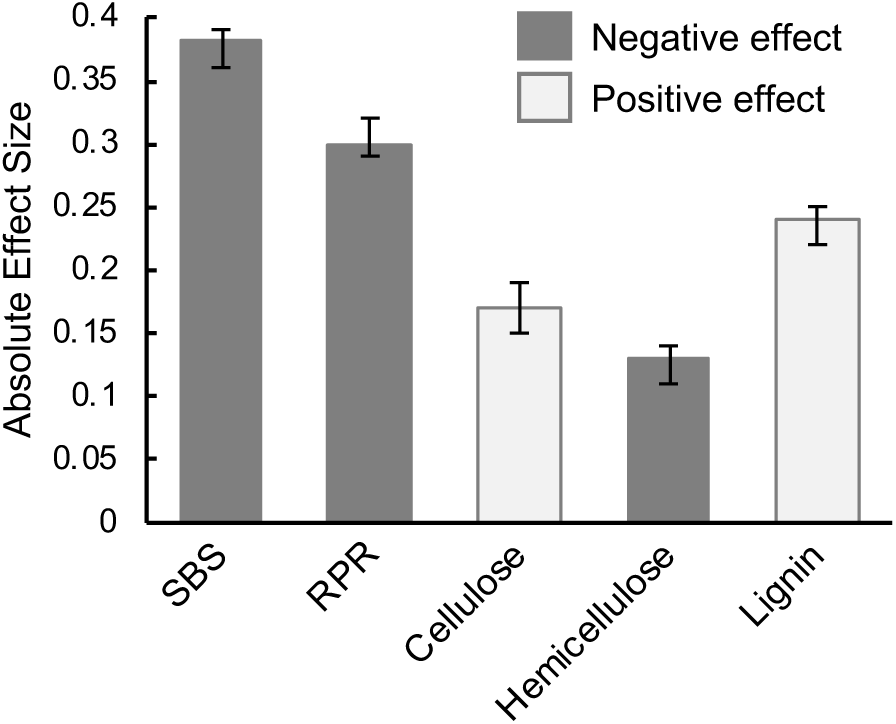
Estimated (absolute) effect sizes for each predictive phenotype with under the marginal models. SBS, stalk bending strength; RPR, rind puncture resistance.

While the marginal models inform about the significance and nature of the association for each of the predictive phenotypes with stalk lodging incidence, a more in-depth analysis is warranted. To this end, we investigated the association of all phenotypes with stalk lodging incidence using a joint model. Effects associated with each predictive phenotype were significant (at the 0.05 significance level) in the joint model (Table 3, Fig. 5). This finding indicates that each predictive phenotype is important in explaining stalk lodging incidence even after accounting for the others. This highlights the fact that each of these predictive phenotypes capture a slightly different aspect of this highly complex trait. In the joint model, stalk bending strength continued to have the largest (in magnitude) effect estimate among all five traits, indicating the strongest association with stalk lodging incidence. Rind puncture resistance also had a negative effect estimate in the joint model, but was substantially less important than stalk bending strength, lignin, and cellulose. Lignin had the largest positive effect estimate. The direction of the estimated effects for cellulose and hemicellulose in the joint model were reversed as compared to those observed in the marginal models.

**Fig. 5.**
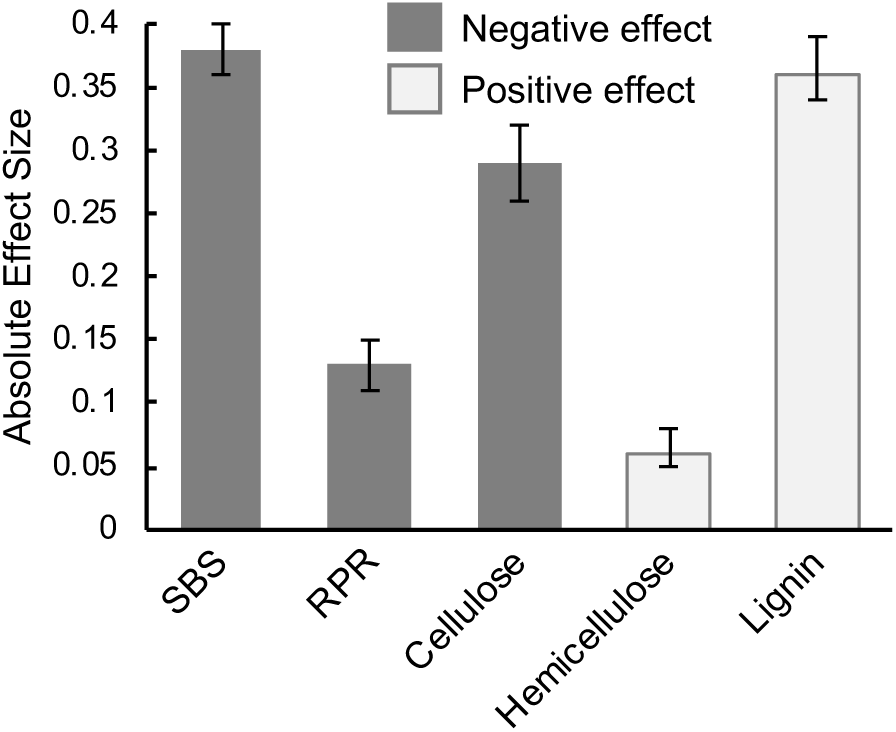
Estimated (absolute) effect sizes for each predictive phenotype under the joint model. SBS, stalk bending strength; RPR, rind puncture resistance.

In summary, while certain aspects of the effect sizes varied between the marginal and joint models, other aspects were consistent. The stalk bending strength had the largest (in magnitude) effect estimate in both modeling approaches and was negatively associated with lodging in both. Similarly, rind puncture resistance was negatively associated with lodging in both approaches. Finally, lignin was positively associated with lodging in both approaches. All things considered, stalk bending strength emerges as the most important predictive phenotype of stalk lodging incidence among those examined in this study.

## 4. Discussion

Determining stalk lodging resistance of individual genotypes has been particularly challenging due to lack of an assessment technique that is quantitative, closely related to lodging, and not confounded by weather events. This study compared the relative importance of multiple quantitative phenotypes with regard to their association with stalk lodging incidence of 47 maize hybrids. We obtained a robust estimate of naturally occurring stalk lodging incidence of each hybrid in an extensive multi-location and multi-year testing initiative. The same hybrids were also evaluated in three distinct environments over two years to produce data on stalk bending strength, rind puncture resistance, and three cell wall metabolites (cellulose, hemicellulose, and lignin). Using Bayesian generalized linear mixed effects models, we demonstrated that, among the phenotypes evaluated, stalk bending strength was the most important predictive phenotype of stalk lodging incidence. We also show that rind puncture resistance is a reasonable predictor of stalk lodging incidence.

Lack of quantitative phenotyping methodologies capable of obtaining accurate measurements of complex plant traits in a cost-effective manner is a major challenge faced by the crop research community. This is evident from relatively slow progress in understanding the genetic architecture and improvement of stalk lodging resistance in maize and related grasses. One of the significant hurdles in the evaluation of a phenotyping method for prediction of stalk lodging resistance is the difficulty in obtaining natural lodging incidence data needed to test the effectiveness of such methods. The occurrence of natural lodging is highly dependent on the environment. As such, genotypes with poor stalk lodging resistance may not exhibit any lodging in studies that include a small number of environments. The G2F initiative is an invaluable source for generating phenotypic data on a large number of diverse locations. The availability of stalk lodging incidence data from 98 distinct environments allowed us to rigorously test the effectiveness and relative importance of the predictive phenotypes presented in this study. Sizeable genotypic differences were observed in stalk lodging incidence. Consistent with a major role of environment, there was substantial variation for stalk lodging incidence across years with 2015 and 2016 being the least and most conducive for lodging, respectively. However, given the geographical diversity of the environments, and therefore the weather pattern experienced by these environments, the overall trend should be treated with caution.

Previous studies have indicated that stalk bending strength is an important determinant of stalk lodging incidence (Gou et al., 2008; Hu et al., 2013). Two unique aspects of the current study are 1) the bending strength data were collected in the field with DARLING as opposed to collected in a laboratory, and 2) the stalk lodging incidence data were obtained from a large number of temporally and geographically distinct environments. To the best of our knowledge, this is the first report wherein a field-based measurement of stalk bending strength has been related to stalk lodging incidence of individual genotypes. Based on the joint model, stalk bending strength measurements produced by DARLING emerged as the most important predictive phenotype of stalk lodging incidence. Given that phenotyping with DARLING can be accomplished rapidly in the field (Cook et al., 2019), this approach may be easily incorporated in breeding programs. The historical lack of an economical and quantitative phenotyping method is responsible for a dearth of information on the genes and genetic elements controlling stalk strength and lodging resistance. To this end, adoption of the DARLING apparatus in the interrogation of existing genetic populations, development of targeted biparental populations, and mutant screens will help elucidate the genetic architecture of this important trait.

Our analyses also demonstrate that rind puncture resistance is a significant predictor of stalk lodging incidence, reinforcing the utility of this approach as a phenotyping strategy. However, rind puncture resistance lacks the inclusiveness of many metabolic and morphological properties of stalks that determine stalk lodging incidence and, therefore lacks sensitivity as compared to bending strength measurements. As such, rind puncture resistance is expected to be effective in early stage breeding trials where larger phenotypic differences are expected among genotypes. However, as the phenotypic differences become progressively smaller due to selection in advanced stages of breeding programs, this method may be less effective. One of the key issues that limits effectiveness of rind puncture resistance as a reliable predictive phenotype is the lack of a standardized testing procedure and associated equipment. Development of a better platform for reproducible rind puncture resistance measurements will likely improve the utility of this approach. When combined with stalk bending strength measurements from DARLING, this method could further enhance the resolution of phenotypic data on stalk lodging incidence.

Composition of stalk cell walls is an important determinant of stalk lodging incidence (Zuber et al., 1980; Appenzeller et al., 2004); however, whether each of these constituents has a positive or negative effect on lodging incidence is debated in the literature. The marginal models showed hemicellulose is negatively associated with lodging incidence while cellulose has a positive association. However, in the joint model that accounts for the relative contribution of all five predictive phenotypes, this trend is reversed. From statistical standpoint, the sign reversal is a well-documented phenomenon that can occur when one moves from a marginal to a joint model. Among other things, this reversal is likely explained by the misspecification of the marginal model resulting from omission of other significant predictive phenotypes in the analysis. That said, the findings from the two analyses are complementary and are aimed at providing a comprehensive overview. From a biological standpoint, this observation highlights the complex relationship between these metabolites in the plant cell wall and strength of the plant organ. This complex relationship is evident from conflicting reports on the role of structural polysaccharides in stalk lodging. For example, while cellulose was proposed to be a positive regulator in one study (Appenzeller et al., 2004), both cellulose and hemicellulose were shown to be of no consequence in another (Ye et al., 2016). The effort to relate the absolute amount of cellulose and hemicellulose with lodging incidence is, perhaps, an over-simplification and the compositional and structural details of these polysaccharides need to be considered. For instance, the extent of crystallinity and degree of polymerization of cellulose affect lodging and, remarkably, arabinose present in hemicellulose negatively affects cellulose crystallinity (Li et al., 2015; Li et al., 2017). A balance between various forms of these two polysaccharides may be of larger consequence in maintaining the cell wall structure and integrity (Sumiyoshi et al., 2013).

Further studies with detailed analyses will shed more light on the association of these polysaccharides with lodging incidence.

Our finding that increased lignification of stalk cell walls is negatively associated with stalk lodging incidence is contrary to some studies which reported a positive impact of this secondary metabolite (Tripathi et al., 2003; Dorairaj et al., 2017; Sun et al., 2018). While this result seems counter-intuitive, there may be a simple explanation. It has been shown that stalk morphology, and in particular the section modulus, is the strongest predictor of stalk bending strength (Robertson et al., 2017), and that lignin is primarily located in the rind tissue. Section modulus is an engineering term used to describe how the mass of an object is distributed about its centroid and is used to calculate the strength and stiffness of three-dimensional structures (Robertson et al., 2017). Since section modulus has been hypothesized to have more influence on stalk strength than tissue properties (Von Forell et al., 2015), stalks with a larger section modulus may have less need for the reinforcing role of lignin. Thus, maize stalks may use lignin strategically, with more lignin required for smaller stalks and less lignin required for larger stalks. The data collected in this study are not sufficient to address this topic fully and further research will be required to investigate this hypothesis. Furthermore, the primary evidence for the role of lignin in the determination of stalk lodging incidence in maize comes from the evaluation of brown midrib (bmr) mutants that show a reduction in lignin content and increased stalk lodging incidence (Miller et al., 1983). However, decrease in lignin content below biologically optimal levels (as in bmr mutants) compromises cell wall strength, alters the plant morphology (e.g., the section modulus), and results in increased stalk lodging incidence. These observations, however, do not necessarily translate into a positive relationship between higher lignification and decreased stalk lodging incidence.

## 5. Conclusions

Stalk bending strength measurements taken on field-grown plants by the novel DARLING apparatus (Cook et al., 2019) emerged as the most important predictive phenotype of stalk lodging incidence. Data also supports the effectiveness of rind puncture resistance as a phenotyping strategy, though this approach lacks the specificity of stalk bending strength measurements. The efficacy of using metabolic composition to predict stalk lodging incidence of genotypes is uncertain based on the available data. While the cell wall components are clearly important for imparting strength to maize stalks, their effect on stalk lodging incidence is unclear. This study paves the way for the adoption of the DARLING apparatus to obtain field-based bending strength measurements in breeding programs and genetics studies to improve lodging resistance in maize and, potentially, in other related grasses.

## Acknowledgements

Authors would like to thank the G2F cooperators and organizers for collecting the lodging incidence data, Naser AlKhalifah for providing raw G2F data, and Matthew Fenton for help with compiling the agronomic details for each of the G2F locations. Funding for this study was provided by National Science Foundation grants #1826715, by the United States Department of Agriculture NIFA Grant #1400973, USDA Hatch project SC-1700520, and National Institutes of Health Grant R01-AI121351.

