## Supplementary figures and images for "Stalk Bending Strength is Strongly Associated with Maize Stalk Lodging Incidence Across Multiple Environments"

### Fig. A.1.

Fig. A.1.

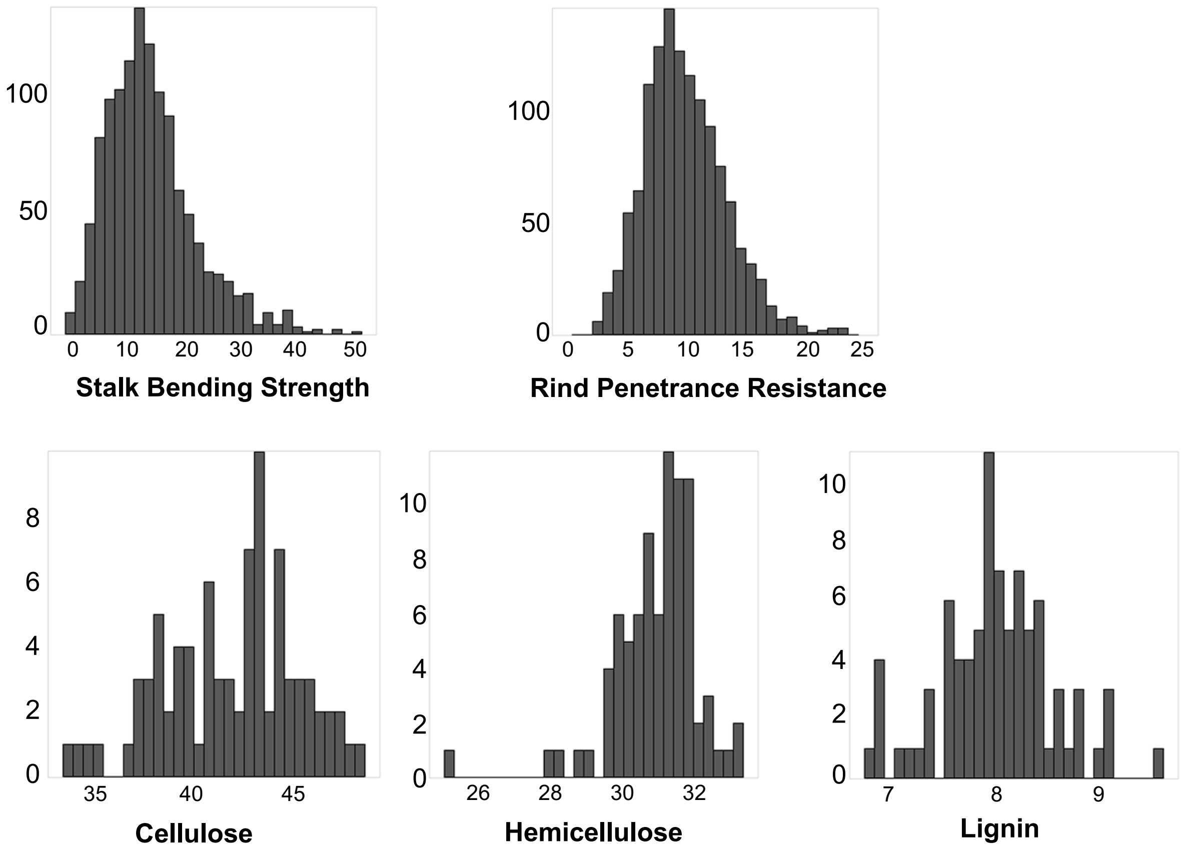


Fig. A.1. Frequency distribution of traits recorded in the study.
