## Supplementary material for "Stalk Bending Strength is Strongly Associated with Maize Stalk Lodging Incidence Across Multiple Environments": Fig. A.2.

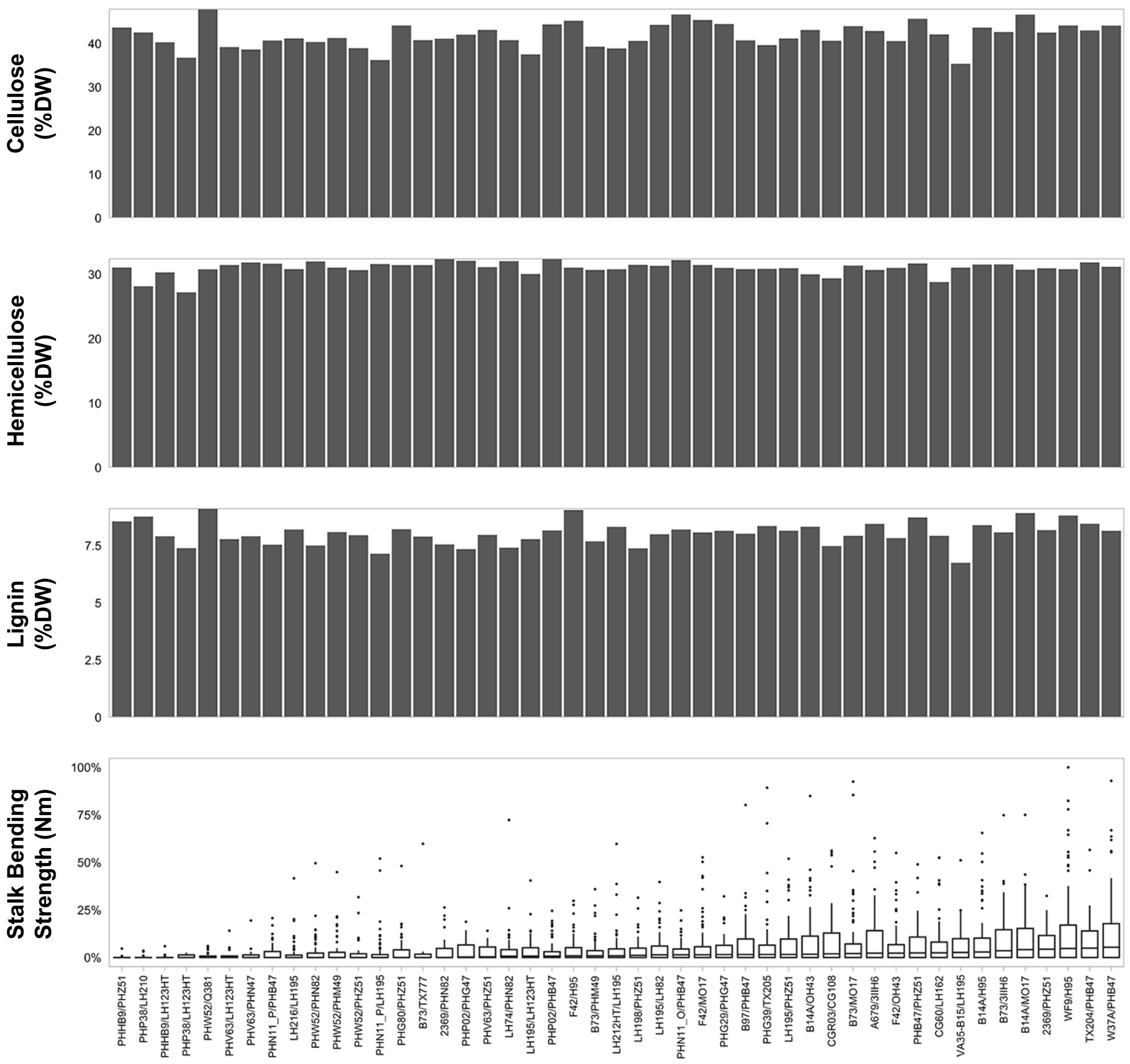


Fig. A.2. Variability in stalk lodging incidence and three metabolic traits captured in the hybrids included in the study. The hybrids are arranged based on increasing median stalk lodging incidence (middle panel). Box plots show the empirical distribution of stalk lodging incidence. In the box plots the lower and upper end of the boxes represent 25th and 75th percentiles of the observed data, respectively; the tips of the vertical lines (whiskers) at the lower and upper end of the boxes represent the 10th and the 90th percentiles of the observed data, respectively; horizontal line within each box denotes the median, and the dots represent outliers. Bar plots represent average content of each of the three metabolites mentioned on y-axis based on two biological replicates. The following hybrid names were shortened: PHN11_PHG47_0251/PHB47 to PHN11_P/PHB47, PHN11_PHW65_0323/LH195 to PHN11_P/LH195, and PHN11_OH43_0001/PHB47 to PHN11_O/PHB47.
