## Supplementary material for "Stalk Bending Strength is Strongly Associated with Maize Stalk Lodging Incidence Across Multiple Environments": Table B.1.

Table B.1. Details of locations and agronomic practices at the Genomes to Fields (G2F) environments used for collection of stalk lodging incidence data

| Location | Year | Longitude | Latitude | Plot Length (m) | Alley Length (m) | Row Spacing (m) | Plant to Plant Spacing (m) | Plants/ plot | Plants/ ha |
|---|---|---|---|---|---|---|---|---|---|
| DEH1 | 2014 | -75.20 | 38.64 | 5.31 | 0.61 | 0.762 | 0.170 | 55.3 | 77,204 |
| GAH1 | 2014 | -83.56 | 31.51 | 6.10 | 1.83 | 0.914 | 0.158 | 54 | 69,197 |
| IAH1a | 2014 | -93.70 | 42.00 | 6.10 | 0.76 | 0.762 | 0.166 | 64.4 | 79,222 |
| IAH1b | 2014 | N/A | N/A | 6.10 | 0.76 | 0.762 | 0.162 | 65.9 | 81,068 |
| IAH1c | 2014 | N/A | N/A | 6.10 | 0.76 | 0.762 | 0.185 | 57.6 | 70,857 |
| IAH2 | 2014 | -94.73 | 42.07 | 6.10 | 0.76 | 0.762 | 0.159 | 67 | 82,421 |
| IAH3 | 2014 | -92.26 | 41.99 | 6.10 | 0.76 | 0.762 | 0.159 | 67 | 82,421 |
| IAH4 | 2014 | -91.49 | 41.20 | 6.10 | 0.76 | 0.762 | 0.156 | 68.2 | 83,897 |
| ILH1 | 2014 | -88.23 | 40.06 | 6.10 | 0.76 | 0.762 | 0.168 | 63.5 | 78,115 |
| INH1 | 2014 | -87.01 | 40.49 | 6.10 | 0.76 | 0.762 | 0.195 | 54.7 | 67,290 |
| MNH1 | 2014 | -93.53 | 44.07 | 8.53 | 0.76 | 0.762 | 0.259 | 60.1 | 50,738 |
| MOH1 | 2014 | -92.21 | 38.90 | 6.10 | 0.76 | 0.914 | 0.204 | 52.3 | 53,615 |
| MOH2 | 2014 | -92.35 | 38.93 | 6.10 | 0.76 | 0.914 | 0.195 | 54.6 | 55,972 |
| NCH1 | 2014 | -77.57 | 35.30 | 4.88 | 0.76 | 0.965 | 0.233 | 35.3 | 44,440 |
| NEH1 | 2014 | -96.66 | 40.83 | 6.10 | 0.76 | 0.762 | 0.168 | 63.5 | 78,115 |
| NEH2 | 2014 | -100.75 | 41.05 | 6.10 | 0.76 | 0.762 | 0.301 | 35.5 | 43,671 |
| NEH3 | 2014 | -102.00 | 41.16 | 6.10 | 0.76 | 0.762 | 0.316 | 33.8 | 41,579 |
| NYH1 | 2014 | -76.65 | 42.73 | 6.40 | 0.76 | 0.762 | 0.186 | 60.5 | 70,402 |
| NYH2 | 2014 | -76.65 | 42.73 | 6.40 | 0.76 | 0.762 | 0.239 | 47.1 | 54,809 |
| ONH1 | 2014 | -80.43 | 43.50 | 6.71 | 0.76 | 0.762 | 0.175 | 68.1 | 75,182 |
| ONH2 | 2014 | -81.88 | 42.45 | 6.10 | 0.76 | 0.762 | 0.212 | 50.3 | 61,877 |
| TXH2 | 2014 | -89.53 | 43.06 | 6.10 | 0.61 | 0.762 | 0.198 | 55.3 | 66,138 |
| WIH1 | 2014 | -89.53 | 43.06 | 7.32 | 0.76 | 0.762 | 0.218 | 60.22 | 60,298 |
| DEH1 | 2015 | -75.47 | 38.63 | 5.30 | 0.61 | 0.762 | 0.169 | 55.4 | 77,444 |
| GAH1 | 2015 | -83.56 | 31.51 | 6.10 | 1.83 | 0.914 | 0.131 | 65.3 | 83,677 |
| ILH1 | 2015 | -88.23 | 40.06 | 6.10 | 0.76 | 0.762 | 0.172 | 62 | 76,270 |
| INH1 | 2015 | -87.00 | 40.48 | 6.10 | 0.76 | 0.762 | 0.203 | 52.5 | 64,583 |
| MNH1 | 2015 | -93.54 | 44.07 | 7.62 | 0.91 | 0.762 | 0.194 | 69.3 | 67,813 |
| MOH1 | 2015 | -92.21 | 38.90 | 6.10 | 0.91 | 0.914 | 0.189 | 54.7 | 57,724 |
| MOH2 | 2015 | -92.21 | 38.90 | 6.10 | 0.91 | 0.914 | 0.184 | 56.2 | 59,307 |
| NCH1 | 2015 | -77.57 | 35.30 | 4.88 | 1.22 | 0.762 | 0.195 | 37.6 | 67,454 |
| NEH2 | 2015 | -100.75 | 41.05 | 6.10 | 0.91 | 0.762 | 0.316 | 32.8 | 41,536 |
| NEH3 | 2015 | -101.99 | 41.16 | 6.10 | 0.76 | 0.762 | 0.298 | 35.8 | 44,040 |
| NYH1 | 2015 | -76.65 | 42.73 | 6.40 | 0.76 | 0.762 | 0.183 | 61.7 | 71,798 |
| NYH2 | 2015 | -76.65 | 42.73 | 6.40 | 1.07 | 0.762 | 0.180 | 59.4 | 73,072 |
| NYH3 | 2015 | -76.66 | 42.72 | 6.40 | 1.07 | 0.762 | 0.165 | 64.7 | 79,591 |
| OHH1 | 2015 | -83.67 | 39.86 | 7.62 | 0.76 | 0.762 | 0.165 | 83.3 | 79,701 |
| ONH1 | 2015 | -80.45 | 43.50 | 6.71 | 0.76 | 0.762 | 0.172 | 69.1 | 76,286 |
| ONH2 | 2015 | -81.88 | 42.45 | 6.10 | 0.76 | 0.762 | 0.170 | 62.9 | 77,377 |
| SDH1 | 2015 | -102.93 | 44.21 | 6.10 | 1.52 | 0.762 | 0.214 | 42.8 | 61,426 |

Table B.1. Details of locations and agronomic practices at the Genomes to Fields (G2F) environments used for collection of stalk lodging incidence data

| Location | Year | Longitude | Latitude | Plot Length (m) | Alley Length (m) | Row Spacing (m) | Plant to Plant Spacing (m) | Plants/ plot | Plants/ ha |
|---|---|---|---|---|---|---|---|---|---|
| TXH2 | 2015 | N/A | N/A | N/A | N/A | N/A | N/A | N/A | N/A |
| WIH1 | 2015 | -89.33 | 43.06 | 7.32 | 0.76 | 0.762 | 0.183 | 71.5 | 71,593 |
| WIH2 | 2015 | -89.33 | 43.33 | 7.32 | 0.76 | 0.762 | 0.187 | 70.1 | 70,191 |
| ARH1 | 2016 | -90.76 | 34.73 | 7.62 | 1.52 | 0.965 | 0.148 | 82.3 | 69,937 |
| ARH2 | 2016 | -90.66 | 35.84 | 4.42 | 1.07 | 0.762 | 0.266 | 25.2 | 49,318 |
| DEH1 | 2016 | -75.45 | 38.65 | 5.30 | 0.71 | 0.762 | 0.165 | 55.5 | 79,300 |
| GAH1 | 2016 | -83.56 | 31.51 | 6.10 | 1.83 | 0.914 | 0.142 | 60 | 76,885 |
| IAH1 | 2016 | -91.49 | 41.20 | 6.10 | 0.76 | 0.762 | 0.171 | 62.4 | 76,762 |
| IAH3 | 2016 | -92.26 | 41.99 | 6.10 | 0.76 | 0.762 | 0.166 | 64.3 | 79,099 |
| IAH4 | 2016 | -93.70 | 42.00 | 6.10 | 0.76 | 0.762 | 0.154 | 69.4 | 85,373 |
| ILH1 | 2016 | -88.23 | 40.06 | 6.34 | 1.02 | 0.762 | 0.161 | 66.1 | 81,469 |
| INH1 | 2016 | -88.99 | 40.48 | 6.10 | 0.76 | 0.762 | 0.164 | 65.2 | 80,207 |
| KSH1 | 2016 | -96.63 | 39.14 | 7.62 | 1.52 | 0.762 | 0.303 | 40.3 | 43,379 |
| KSH2 | 2016 | -100.78 | 37.81 | 6.10 | 0.76 | 0.762 | 0.368 | 29 | 35,675 |
| KSH3 | 2016 | -100.78 | 37.81 | 6.10 | 0.76 | 0.762 | 0.374 | 28.5 | 35,060 |
| MIH1 | 2016 | -84.50 | 42.69 | 7.62 | 0.91 | 0.762 | N/A | N/A | N/A |
| MOH1 | 2016 | -92.21 | 38.90 | 6.10 | 0.91 | 0.762 | 0.147 | 70.3 | 89,024 |
| NCH1 | 2016 | -77.57 | 35.30 | 4.88 | 1.22 | 0.762 | 0.207 | 35.4 | 63,507 |
| NYH1 | 2016 | -76.65 | 42.73 | 6.40 | 0.91 | 0.762 | 0.170 | 64.6 | 77,261 |
| NYH2 | 2016 | -76.66 | 42.73 | 6.40 | 1.07 | 0.762 | 0.333 | 32 | 39,365 |
| NYH3 | 2016 | -76.66 | 42.73 | 6.40 | 1.07 | 0.762 | 0.171 | 62.3 | 76,639 |
| OHH1 | 2016 | -83.68 | 39.86 | 7.62 | 0.76 | 0.762 | 0.161 | 85.2 | 81,519 |
| ONH1 | 2016 | -80.45 | 43.50 | 5.94 | 0.76 | 0.762 | 0.144 | 72 | 91,177 |
| ONH2 | 2016 | -81.88 | 42.45 | 5.33 | 0.76 | 0.762 | 0.145 | 63.2 | 90,704 |
| SCH1 | 2016 | -82.74 | 34.62 | 6.10 | 1.52 | 0.762 | 0.195 | 46.8 | 67,167 |
| WIH1 | 2016 | -89.53 | 43.06 | 7.32 | 0.61 | 0.762 | 0.208 | 64.6 | 63,214 |
| WIH2 | 2016 | -89.34 | 43.33 | 7.32 | 0.61 | 0.762 | 0.188 | 71.3 | 69,770 |
| ARH1 | 2017 | -90.76 | 34.73 | 7.62 | 1.52 | 0.965 | 0.157 | 77.8 | 66,113 |
| ARH2 | 2017 | -90.08 | 35.67 | 7.62 | 1.52 | 0.965 | 0.174 | 69.9 | 59,400 |
| COH1 | 2017 | -105.00 | 40.65 | 5.33 | 0.76 | 0.762 | 0.164 | 55.6 | 79,796 |
| DEH1 | 2017 | -75.43 | 38.67 | 5.31 | 0.61 | 0.762 | 0.164 | 57.4 | 80,136 |
| GAH1 | 2017 | -83.56 | 31.51 | 6.10 | 1.83 | 0.914 | 0.240 | 35.6 | 45,618 |
| GAH2 | 2017 | -83.30 | 33.73 | 7.62 | 0.13 | 0.762 | 0.344 | 43.6 | 38,181 |
| IAH1 | 2017 | -91.50 | 41.20 | 6.10 | 0.76 | 0.762 | 0.170 | 62.9 | 77,377 |
| IAH2 | 2017 | -94.72 | 42.06 | 6.10 | 0.76 | 0.762 | 0.158 | 67.6 | 83,159 |
| IAH3 | 2017 | -92.24 | 41.98 | 6.10 | 0.76 | 0.762 | 0.174 | 61.4 | 75,532 |
| IAH4 | 2017 | -93.69 | 41.99 | 6.10 | 0.76 | 0.762 | 0.157 | 67.9 | 83,528 |
| ILH1 | 2017 | -89.38 | 43.07 | 6.33 | 1.01 | 0.762 | 0.191 | 55.7 | 68,700 |
| INH1 | 2017 | -89.38 | 43.07 | 6.10 | 0.76 | 0.762 | 0.178 | 59.8 | 73,564 |
| MIH1 | 2017 | -84.49 | 42.68 | 7.62 | 0.76 | 0.762 | 0.172 | 79.7 | 76,256 |

Table B.1. Details of locations and agronomic practices at the Genomes to Fields (G2F) environments used for collection of stalk lodging incidence data

| Location | Year | Longitude | Latitude | Plot Length (m) | Alley Length (m) | Row Spacing (m) | Plant to Plant Spacing (m) | Plants/ plot | Plants/ ha |
|---|---|---|---|---|---|---|---|---|---|
| MNH1 | 2017 | -89.38 | 43.07 | 7.62 | 0.91 | 0.762 | 0.198 | 67.8 | 66,345 |
| MOH1 | 2017 | -92.21 | 38.89 | 6.10 | 0.91 | 0.762 | 0.173 | 59.9 | 75,854 |
| NCH1 | 2017 | -77.57 | 65.30 | 4.88 | 1.22 | 0.762 | 0.196 | 37.3 | 66,916 |
| NEH3 | 2017 | -89.38 | 43.07 | 5.33 | 0.76 | 0.762 | 0.260 | 35.2 | 50,519 |
| NEH4 | 2017 | -89.38 | 43.07 | 5.33 | 0.76 | 0.762 | 0.278 | 32.9 | 47,218 |
| NYH1 | 2017 | -89.38 | 43.07 | 5.33 | 0.76 | 0.762 | 0.163 | 56 | 80,371 |
| NYH2 | 2017 | -76.65 | 42.73 | 6.40 | 1.07 | 0.762 | 0.183 | 58.3 | 71,718 |
| NYH3 | 2017 | -76.65 | 42.73 | 6.40 | 1.07 | 0.762 | 0.173 | 61.6 | 75,778 |
| OHH1 | 2017 | -83.67 | 39.86 | 7.62 | 0.76 | 0.762 | 0.209 | 65.6 | 62,766 |
| ONH1 | 2017 | -80.43 | 43.50 | 6.71 | 0.76 | 0.762 | 0.164 | 72.5 | 80,039 |
| ONH2 | 2017 | -89.38 | 43.07 | 6.10 | 0.76 | 0.762 | 0.160 | 66.5 | 81,806 |
| SCH1 | 2017 | -89.38 | 43.07 | 7.62 | 1.22 | 0.635 | 0.311 | 41.2 | 50,683 |
| TXH1-Dry | 2017 | -96.43 | 30.55 | 7.62 | 1.22 | 0.635 | 0.211 | 60.6 | 74,548 |
| TXH1-Early | 2017 | -96.43 | 30.55 | 7.62 | 1.22 | 0.635 | 0.206 | 62 | 76,270 |
| TXH1-Late | 2017 | -96.43 | 30.55 | 7.62 | 1.22 | 0.635 | 0.204 | 62.9 | 77,377 |
| TXH2 | 2017 | -89.38 | 43.07 | 5.49 | 0.61 | 1.016 | 0.137 | 71.1 | 71,748 |
| WIH1 | 2017 | -89.53 | 43.06 | 5.49 | 0.76 | 0.762 | 0.146 | 64.8 | 90,000 |
| WIH2 | 2017 | -89.34 | 43.32 | 5.49 | 0.76 | 0.762 | 0.137 | 68.8 | 95,556 |
